# Role of Lamin A/C on dendritic cell function in antiviral immunity

**DOI:** 10.1101/2024.05.14.593747

**Authors:** Beatriz Herrero-Fernandez, Raquel Gómez Bris, Marina Ortega Zapero, Angela Saez, Salvador Iborra, Virginia Zorita, Ana Quintas, Enrique Vazquez, Ana Dopazo, Francisco Sánchez-Madrid, Silvia Magdalena Arribas, Jose Maria Gonzalez Granado

## Abstract

Dendritic cells (DCs) play a crucial role in orchestrating immune responses, particularly in promoting IFNγ-producing-CD8 cytotoxic T lymphocytes (CTLs) and IFNγ-producing -CD4 T helper 1 (Th1) cells, which are essential for defending against viral infections. Additionally, the nuclear envelope protein lamin A/C has been implicated in T cell immunity. Nevertheless, the intricate interplay between innate and adaptive immunity in response to viral infections, particularly the role of lamin A/C in DC functions within this context, remains poorly understood. In this study, we demonstrate that mice lacking lamin A/C in myeloid LysM promoter-expressing cells exhibit a reduced capacity to induce Th1 and CD8 CTL responses, leading to impaired clearance of acute primary *Vaccinia virus* (VACV) infection. Remarkably, *in vitro*-generated granulocyte macrophage colony-stimulating factor bone marrow-derived DCs (GM-CSF BMDCs) show high levels of lamin A/C. Lamin A/C absence on GM-CSF BMDCs does not affect the expression of costimulatory molecules on the cell membrane but it reduces the cellular ability to form immunological synapses with naïve CD4 T cells. Lamin A/C deletion induces alterations in NFκB nuclear localization, thereby influencing NFκB-dependent transcription. Furthermore, lamin A/C ablation modifies the epigenetic signature of BMDCs, predisposing these cells to mount a less effective antiviral response upon TLR stimulation. This study highlights the critical role of DCs in interacting with CD4 T cells during antiviral responses and elucidates the molecular mechanisms through which lamin A/C modulates DC function via epigenetic and transcriptional regulation.

## Introduction

Viruses exploit various entry routes to penetrate body barriers, infect diverse cell types, and adapt to evade immune surveillance. Eradicating rapidly replicating viruses effectively requires a multifaceted immune response orchestrated by different cellular entities targeting various viral components. Upon infection, the innate immune system promptly mobilizes to counteract viral dissemination, thus triggering the adaptive arm to initiate a specific immune defense, ultimately resulting in viral clearance [1].

The immunization with the *Vaccinia virus* (VACV), a linear double-stranded DNA virus, has shown efficacy against *Variola major*, contributing significantly to the global eradication of smallpox [2]. Notably, the priming, activation, proliferation, and differentiation of T cells in response to VACV rely on professional antigen-presenting cells (APCs), particularly dendritic cells (DCs) [3].

DCs can be categorized into distinct groups based on phenotype, function, and development [4,5], including plasmacytoid DCs (pDCs), conventional DCs (cDCs) and, monocyte-derived DCs (moDCs). Under homeostasis, immature DCs are distributed throughout the body and at microbial entry points, serving as sentinels for danger signals. Common DC progenitors are the primary source of DCs. However, in mice, inflammation at infected sites triggers the recruitment of Ly6C^high^ monocytes from the bloodstream [6]. Upon arrival, these monocytes differentiate into moDCs [7] in the presence of granulocyte-macrophage colony-stimulating factor (GM-CSF) [8], which is produced by activated T cells and other cell types in the inflamed tissue [8]. In addition to tissue-resident DCs, moDCs initiate additional rounds of T cell priming [9].

Upon viral encounter in mice and humans, both peripheral tissue-resident DCs and moDCs recognize pathogens via germ-line encoded pattern recognition receptors (PRRs), including Toll-like receptors (TLRs), RIG-I-like receptors (RLRs) and cytoplasmic DNA sensors [10]. Among these, TLR signaling cascades, bifurcated into MyD88-dependent pathway and TRIF-dependent pathway. TLR4 signaling depends on both adaptor molecules, TRIF and MyD88, while TLR2 depends on MyD88 [11]. Upon detection of PAMPs, the Toll/Interleukin-1 receptor (TIR) domain-containing adaptor proteins, MyD88 and/or TRIF, are recruited to the TLRs. This initiation triggers signaling pathways that lead to the activation of NF-κB, interferon regulatory factors (IRFs), and mitogen-activated protein kinase (MAPK) cascades. These pathways ultimately induce the upregulation of costimulatory molecules (CD40, CD80, and CD86), inflammatory cytokines (such as IL-12, IL-6, and TNF), chemokines (RANTES, IP-10, ENA78, etc.), and type I interferons (IFNs) [12–14].

Maturation of both tissue-resident DCs and moDCs is imperative for effective priming of naïve CD8 and CD4 T cells, thereby initiating adaptive immunity [15] via major histocompatibility complex class I/II (MHCI/II) molecules to CD8 and CD4 T cells, respectively [16]. Upon antigen recognition, CD8 T cells differentiate into cytotoxic T lymphocytes (CD8 CTLs), which exert precise antiviral effects via cytokine secretion and release of cytotoxic granules [17,18]. Similarly, CD4 T cells differentiate into T helper 1 cells (Th1), orchestrating antiviral responses through IFNγ production [18], engaging in direct cytotoxic mechanisms [18–20] and assist in B cell-mediated viral clearance via affinity maturation and antibody class switching [21]. Additionally, they contribute to CD8 T cell expansion and maintenance and memory against the virus [22,23].

In mammalian cells, the nuclear envelope is composed of the nuclear pore complex, embedded in both the outer and inner nuclear membranes, along with the nuclear lamina. The nuclear lamina is composed of two main components, A- and B-type lamins, which contribute to the mechanical stability of the nucleus and regulate nuclear positioning, chromatin structure, the organization of the nuclear pore complex, the behavior of the nuclear envelope during mitosis, DNA replication, DNA damage responses, cell-cycle progression, cell differentiation, cell polarization during migration, and transcription [24,25]. Our previous research has demonstrated the pivotal role of lamin A/C in enhancing T cell activation and promoting Th1 responses, which are crucial for effective viral clearance.[26,27] [20]. Furthermore, the absence of lamin A/C in T cells promotes their differentiation towards a Treg phenotype [28,29]. Despite extensive information on the importance of lamin A/C in T cell immunity, its significance in DC functions, particularly in orchestrating the interplay between innate and adaptive immune responses to viral infections, remains an area that warrants further investigation.

This study provides valuable insights into the role of DCs in interacting with CD4 T cells in the antiviral response and the underlying molecular mechanisms controlled by lamin A/C, involving epigenetic and transcriptional regulation in DCs.

## Results

### Reduced lamin A/C levels in the myeloid compartment partially impede the response to VACV by affecting T cell-mediated stimulation

The importance of lamin A/C within the myeloid compartment and the functional role of in *vitro* generated GM-CSF BMDCs were evaluated through various *in vivo* and *in vitro* infection and inflammation approaches.

To selectively downregulate lamin A/C expression in specific myeloid cells, including moDCs and *in vitro* generated-BMDCs, we established LysM-cre^+/-^ Lmna^flox/flox^ mice referred to as LysM-Lmna^-/-^ or Lmna^-/-^ hereafter. LysM-Lmna^-/-^ mice exhibited physical characteristics and body weight comparable to wild-type (WT) mice with no discernible differences in the size, weight, and morphology of various immune system organs, such as the spleen or thymus, when compared to WT mice (described in [30]). Immunofluorescence analysis of cell populations in homeostasis in organs such as the spleen, mesenteric lymph nodes (mLN), lung, or bone marrow (BM) revealed no notable differences between genotypes, except for a reduction in the percentage of alveolar macrophages in adult LysM-Lmna^-/-^ animals within the studied populations [30].

To explore the impact of lamin A/C in the myeloid compartment in the response to the virus, mice were infected with VACV, known for inducting robust Th1 immune response in humans [31] and in mice [1,20,32]. Given the role of CD4 T cells in regulating the primary CD8 T cell response to VACV infection depending on the route of infection and viral dose [33,34], various administration routes were employed including subcutaneous (s.c.), inoculation in the footpad, intraperitoneal injection (i.p.), or via skin scarification (s.s.) in the tail vein (Figure 1).

**Figure 1.**
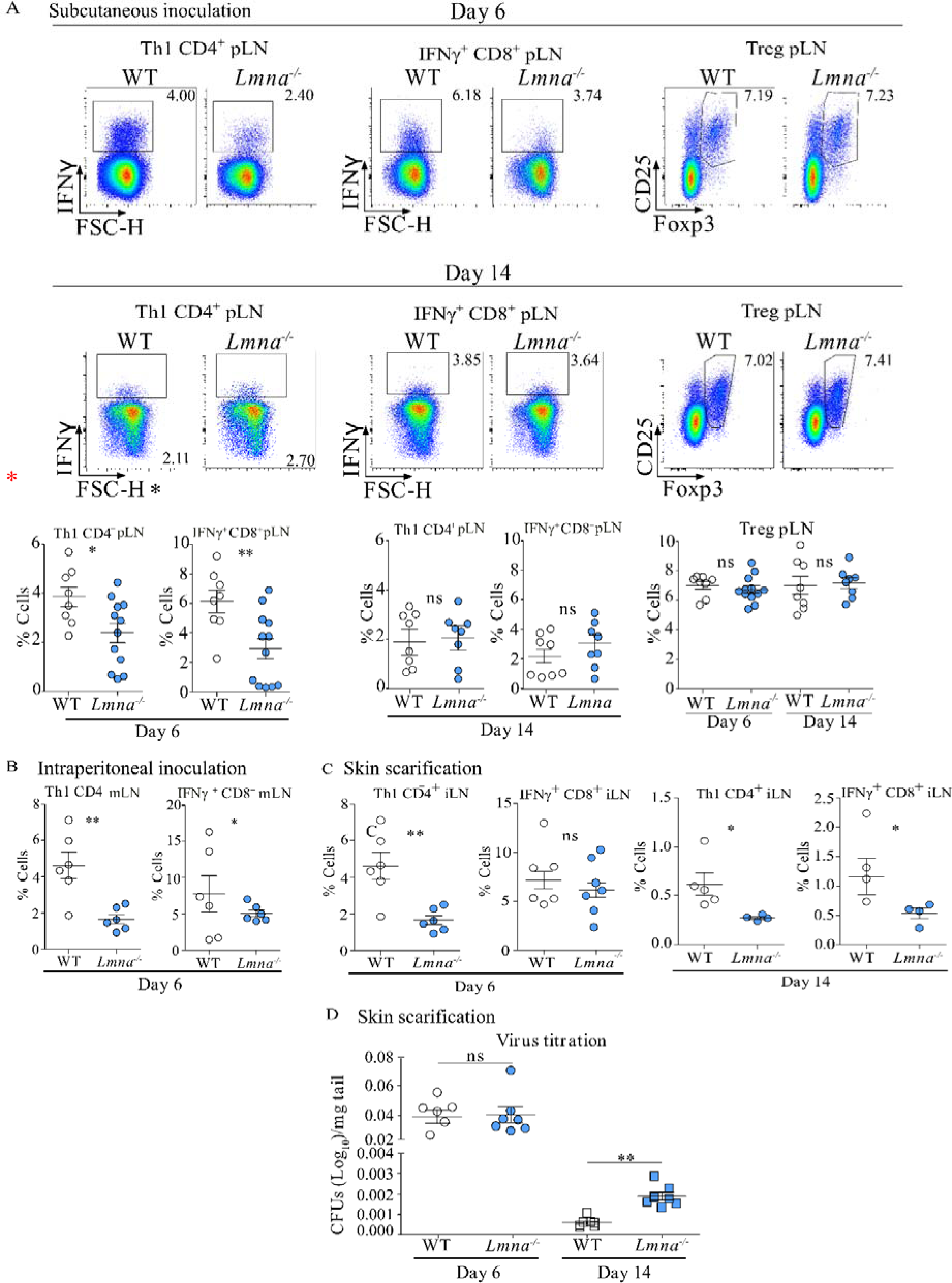
Lamin A/C modulation in myeloid compartment influences vaccinia virus susceptibility and protection across infection routes. (A) Subcutaneous inoculation (s.c.) with 10^5^ p.f.u. of VACV was administered to both WT and LysM-Lmna^-/-^ mice. Flow cytometry analysis of IFNγ-producing CD4 and CD8 T cells, as well as CD25^+^Foxp3^+^ Treg cells, was conducted in popliteal lymph nodes (pLN) at 6 and 14 days post-infection. The percentage of Th1 cells, CD8 CTL T cells, and Treg cells was assessed. The data represent means ± SEM from at least 8 mice per group, obtained from 3 independently conducted experiments, and were analyzed using unpaired Student’s t-test between genotypes at each time point. Statistical significance is denoted by asterisks (*P < 0.05; **P < 0.01; ns, not significant). (B) Intraperitoneal inoculation (i.p.) with 10^6^ p.f.u. of VACV was performed on both WT and LysM-Lmna^-/-^ mice. Flow cytometry analysis of IFNγ-producing CD4 and CD8 T cells was conducted in mesenteric lymph nodes (mLN) 6 days post-infection. The percentage of Th1 and CD8 CTL cells was determined. Data represent means ± SEM of at least 6 mice per group, derived from 2 independently conducted experiments, and were analyzed by unpaired Student’s t-test. Statistical significance is indicated by asterisks (*P < 0.05; **P < 0.01). (C) Infection with 10^5^ p.f.u. of VACV through skin scarification (s.s.) in the tail was performed on WT and LysM-Lmna^-/-^ mice. Flow cytometry analysis of the percentages of IFNγ-producing CD4 and CD8 T cells was conducted in extracted inguinal lymph nodes (iLN) at 6 and 14 days post-infection. (D) Viral load measurement in the tail skin was performed at 6 and 14 days post-infection. The data represent means ± SEM of at least 4 mice per group, derived from 2 independently conducted experiments, and were analyzed by unpaired Student’s t-test. Statistical significance is denoted by asterisks (*P < 0.05; **P < 0.01; ns, not significant).

In the initial infection experiments, both WT and LysM-Lmna-/- mice were administered VACV via the footpad (s.c.). Subsequently, at 6 and 14 days post-infection, popliteal lymph nodes (pLN) were harvested for the analysis of T cell populations (Figure 1A). The frequencies of Treg cells and IFNγ-producing CD4 and CD8 T cells were assessed using flow cytometry. Only at the 6-day post-infection time point were differences observed between genotypes in the percentage of IFNγ-producing CD4 and CD8 T cells. No disparities were noted at the 14-day time point or in the percentage of Treg cells at either of the two time points analyzed.

Further experiments were conducted by inoculating WT and Lmna^-/-^ mice i.p. with VACV (Figure 1B). Six days later, the frequency of IFNγ-producing CD4 and CD8 T cells was analyzed in mLN using flow cytometry. Interestingly, in the mLN of LysM-Lmna^-/-^ mice, there was a reduced percentage of both IFNγ^+^ CD4 and CD8 T cells at 6 days.

Similar outcomes were observed in mice administered VACV through s.s. in the tail vein (Figure 1C). In this context, the percentage of IFNγ^+^ CD4 and CD8 T cells in the inguinal lymph nodes (iLN) was evaluated at two distinct time points: 6 and 14 days. While no statistically significant differences were noted at 6 days in IFNγ^+^ CD8 T cells, there was a significant reduction in IFNγ-producing CD4 T cells (Figure 1C). By 14 days post-infection, a reduced number of both IFNγ-producing CD4 and CD8 T cells was observed in Lmna^-/-^ mice (Figure 1C). This trend correlated with viral titers at the infection site. We observed that VACV levels in the skin of infected mice were higher at 6 days than at 14 days post-infection through a s.s. route in WT mice. Interestingly, the virus levels at 6 days did not differ between WT and Lmna^-/-^ mice. However, in WT mice, there was a subsequent reduction in virus levels in the skin by 14 days. This reduction was significantly less pronounced in Lmna^-/-^ mice, resulting in higher virus levels in the skin at 14 days in Lmna^-/-^ mice compared to WT mice (Figure 1D). These findings underscore the importance of lamin A/C in myeloid cells for efficient virus clearance, particularly by influencing the stimulation of the adaptive immune system, specifically the Th1-cell-mediated and CD8 cytotoxic T lymphocyte (CTL) responses.

### Deficiency of Lamin A/C in BMDCs influences impaired T cell responses across maturation and stimulation environments

To resemble conditions similar to *in vivo* mo-DCs, bone marrow cells were cultured with GM-CSF [35]. The influence of lamin A/C on GM-CSF BMDCs (referred as BMDCs hereafter) in T cell activation, proliferation, and differentiation was then assessed under varied conditions. In a setup like VACV infection, BMDCs were exposed to irradiated VACV and then loaded with ovalbumin (OVA) peptide OVA_323–339_ (OVA-OT-IIp). These cells were co-cultured with naïve CD4 OT-II T cells, which express a TCR recognizing the OVA-OT-IIp, for 24h or 48h for activation analysis and for 6 days for proliferation and differentiation towards the Th1 phenotype analysis, respectively. Lmna^-/-^-BMDCs co-cultures consistently showed a lower percentage of activated CD4 T cells at all observed time points, shown by decreased membrane expression of CD69 and CD25 in CD4 T cells (Figure 2A). Additionally, these CD4 T cells displayed reduced proliferation capacity and diminished differentiation toward the Th1 phenotype, reflected in a reduction in the percentage of cells expressing IFNγ (Figure 2A).

**Figure 2:**
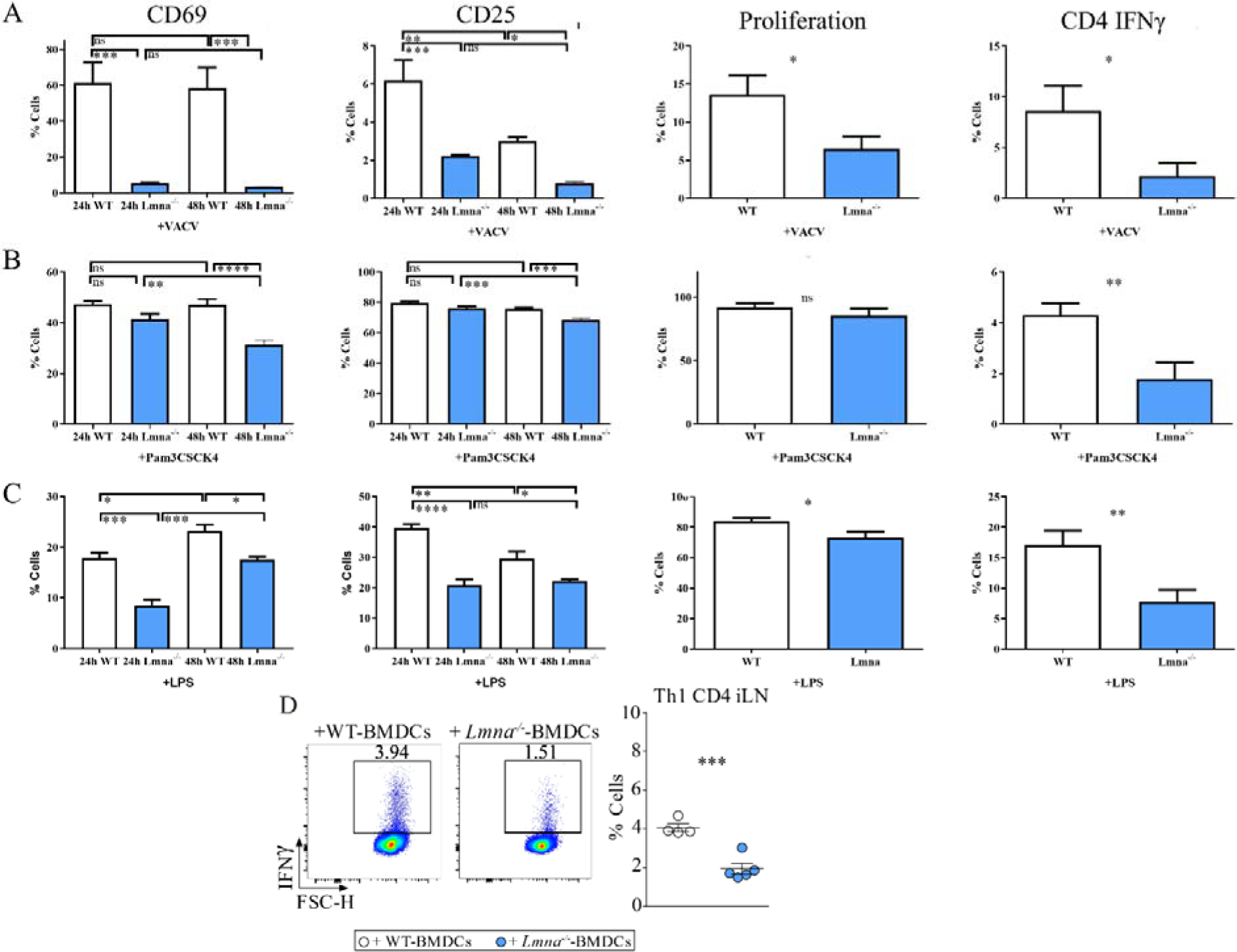
Lamin A/C deficiency in matured GM-CSF-BMDCs reduces CD4 T cell activation, proliferation and Th1 differentiation *in vitro* and *in vivo*. Flow cytometry was employed to assess the percentage of activated CD4 T cells, specifically gated on CD25^+^ or CD69^+^ cells; Th1-differentiated cells, specifically gated on IFNγ+ cells; and proliferating CellTrace Violet^+^ T cells. Maturation of GM-CSF-derived WT- and LysM-Lmna^-/-^-BMDCs was undertaken with (A) sonicated VACV extract, (B) Pam3CSK4, or (C) lipopolysaccharide (LPS), and, in all conditions, pulsed with _OVA323–339_ cognate OT-IIp. (D) WT- or Lmna^-/-^-BMDCs, matured with LPS and pulsed with the OVA_323–339_ cognate OT-II peptide, were s.c. inoculated into CD45.1 OT-II recipient mice. Six days later, pLNs were extracted, and IFNγ-producing CD4 T cells were analyzed by flow cytometry. The data represent means ± SEM and are presented from a representative experiment out of 3 (A and B, n=3) or out of 4 (C, n=4) or out of 2 (D, n=4). Activation data were analyzed using One Way ANOVA and Bonferroni post-test, and proliferation and Th1 differentiation were assessed by unpaired Student’s t-test. Statistical significance was denoted by asterisks (*P < 0.05; **P < 0.01; ***P < 0.001; ns, not significant).

In the immune system, the detection of viruses by DCs/APCs is not solely dependent on nucleic acid–specific receptors. Instead, some TLRs, such as TLR2 and TLR4, which do not recognize nucleic acids, play a crucial role in recognizing various viruses. This recognition is essential for distinguishing between different classes of pathogens to facilitate an appropriate immune response [36].

Therefore, given that VACV influences DCs, in part through TLR2 [37,38] and TLR4 [39,40], we stimulated BMDCs with TLR2 ligand Pam3CSK4 and TLR4 ligand LPS. Subsequently, BMDCs were pulsed with OVA-OT-IIp and co-cultured with naïve CD4 OT-II T cells for 24h or 48h, and for 6 days. Upon TLR2 stimulation (Figure 2B), Lmna^-/-^-BMDCs consistently induced a lower percentage of activated CD4 T cells at 48h. Furthermore, these CD4 T cells displayed a diminished capacity to differentiate toward the Th1 phenotype, with no observed differences in proliferation rates between genotypes. In the context of LPS-induced maturation for TLR4 stimulation, Lmna^-/-^-BMDCs exhibited a lower percentage of activated CD4 T cells at 24 and 48h. Importantly, these CD4 T cells also showed reduced proliferation and differentiation toward the Th1 phenotype (Figure 2C).

To assess the stimulatory potential of these BMDCs within a more physiological context, *in vivo* adoptive transfer experiments were conducted. CD45.1 OT-II mice received s.c. injection in footpad with BMDCs matured with LPS and loaded with OVA-OT-IIp. Six days post-BMDCs injection, draining pLN were harvested, and Th1 differentiation was evaluated. Interestingly, recipient mice inoculated with Lmna^-/-^-BMDCs exhibited lower Th1 differentiation, as indicated by a reduced percentage of IFNγ^+^ CD4 T cells (Figure 2D). These results indicate that the modulation of lamin A/C in BMDCs significantly impacts T cell responses, leading to decreased activation, proliferation, and differentiation, particularly under various maturation and stimulation conditions. This influence is probably mediated through mechanisms associated with TLR2 or TLR4 signaling pathways.

### Lamin A/C in BMDCs controls the formation of immune synapse

Since some studies suggest that TLR2 might recognize virus preparations *in vitro* but plays a minor role in preventing VACV dissemination following systemic infection with large viral doses [41], and considering that TLR2 activation by VACV might be linked to Th2 conversion and robust humoral responses [42], which contrasts with the Th1 and CTL response observed in our *in vivo* studies associated with LPS stimulation [42], we explore the impact of lamin A/C on TLR4 signaling by stimulating with LPS. During antigen recognition, T cells and DCs establish communication through the immune synapse (IS), which is characterized by the precise redistribution and interaction of the TCR/CD3 at the T cell side and the MHC-II at the APC side, along with costimulatory and adhesion molecules on both sides [43]. Lamin A/C is crucial for TCR-dependent signaling and cytoskeletal reorganization at the T cell side [26,27,29]. To investigate this on the APC side, time-lapse confocal microscopy experiments were conducted. Analysis of the interaction between OT-II CD4+ T cells and LPS-matured OVA OT-IIp-loaded WT- or Lmna^-/-^-BMDCs revealed that CD4 T cells exhibited higher speed and reduced interaction time when co-cultured with Lmna^-/-^-BMDCs (Figure 3A). This suggests d a reduced ability of LPS-matured Lmna^-/-^-BMDCs to form stable conjugates, which may contribute to reduced activation, proliferation, and differentiation of T cells toward a Th1 phenotype [26,27].

**Figure 3:**
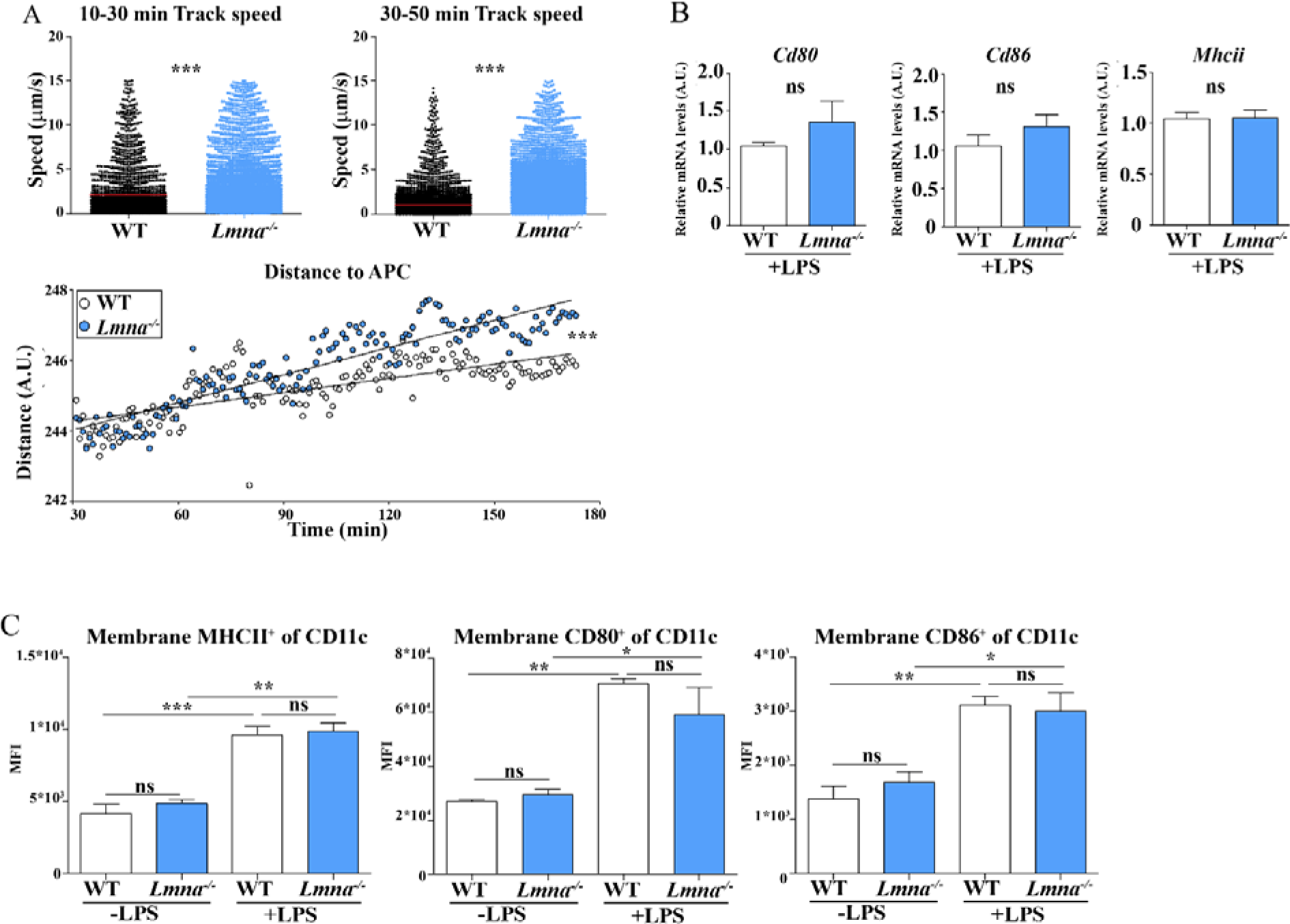
Reduced lamin A/C levels in matured GM-CSF-BMDCs diminish the formation of conjugates with T cells, without affecting the gene expression or membrane localization of MHC-II and costimulatory molecules. (A) WT- and Lmna^-/-^-BMDCs, matured with LPS and pulsed with OVA323–339 cognate OT-IIp, were co-cultured with OT-II CD4 T cells in a 1:1 ratio. Prior to co-culture, BMDCs were stained with DiL, and OT-II CD4 T cells were pre-labeled with CMAC. The study focused on evaluating the speed (µm/s) and distance (A.U.) at different time points of cell-cell contacts. The presented data is from a representative experiment out of 2, and each symbol in the graphs represents an individual cell-cell conjugate (n ≥ 250). The analysis was conducted using unpaired Student’s t-test for the speed and linear regression for the distance. Statistical significance for the distance is indicated by **P < 0.001. A.U. denotes arbitrary units. (B) RT-qPCR analysis was performed on indicated genes from WT- and Lmna^-/-^-BMDCs, both stimulated with LPS for 24 hours. The data, obtained from three independent experiments with 3 samples each, revealed no significant differences in the expression of Mhc-ii and costimulatory molecules genes between the two genotypes (n = 3). The analysis employed unpaired Student’s t-test, and the housekeeping genes *Ywhaz,* β*-actin*, and *18s* were used for normalization. The data represent means ± SEM. (ns, not significant). (C) The membrane levels of MHC-II, CD80, and CD86 were evaluated on CD11c+ cells derived from WT and Lmna^-/-^-BMDCs, either matured or not with LPS. The geometric mean fluorescence intensity (MFI) of MHC-II+, CD80+, and CD86+ markers was analyzed. The data, representing means ± SEM from three independently conducted experiments (n = 3), underwent statistical analysis using one-way ANOVA with Bonferroni’s multiple comparison test. Significance levels are denoted by asterisks (*P < 0.05; **P < 0.01; ***P < 0.001; ns, not significant).

Given the observed disparities in the ability of BMDCs to form T cell conjugates, we investigated whether BMDCs exhibited distinct expression levels of MHC-II and costimulatory molecules, including *Cd80*, and *Cd86*. mRNA analysis showed no discernible differences between genotypes in LPS-matured BMDCs. (Figure 3B). The maturation process of BMDCs involves an upregulation of membrane expression levels of MHC-II, CD80, and CD86, which enhances T cell stimulation [44]. To assess their membrane expression by flow cytometry, we examined both LPS-untreated and LPS-treated matured BMDCs (Figure 3C). As anticipated, there was a noticeable increase in the levels of costimulatory molecules associated with maturation. It is noteworthy that no significant differences were observed in the expression levels between genotypes in unmatured and LPS-matured conditions.

### Lamin A/C in BMDCs promotes nuclear localization of NF-**κ**B

TLR4 signaling stimulates the expression of Nuclear factor kappa-light-chain-enhancer of activated B cells (NF-κB)-related genes expression [45]. NF-κB plays a crucial role in regulating DC development, survival, and cytokine production [46]. As a master nuclear transcription factor for genes involved in inflammatory responses, RelA/p65, together with p50, constitutes the most common NF-κB subunits involved in the classical NF-κB pathway. NF-κB activity is tightly controlled by inhibitors, such as IκBα, which sequesters p65/p50 dimers in the cytoplasm. Activation of various signaling cascades leads to increased IκB kinase (IKK) activities, resulting in IκBα phosphorylation and subsequent degradation. This allows the release of homo- or hetero-dimers of NF-κB, enabling their translocation into the nucleus, where they activate gene transcription [47].

Given that previous studies have indicated lamin A/C role in enhancing the nuclear translocation of RelA/p65 and subsequently increasing NF-κB activity, leading to elevated inflammatory gene expression in macrophages [48], we sought to investigate whether a similar phenomenon occurs in our BMDC model. To explore this, WT- or Lmna^-/-^-BMDCs were either treated or left untreated with LPS for 24 hours, and confocal images of lamin A/C and NF-κB staining were acquired. The nucleus was identified by DAPI staining (Figure 4). The analysis of NF-κB subunit, RelA, translocation to the nucleus is a well-established method for determining NF-κB activation [48]. Upon LPS addition, nuclear levels and the nuclear/cytoplasmic ratio of NF-κB significantly increased in WT BMDCs, without affecting total cellular NF-κB levels. However, Lmna^-/-^ BMDCs did not exhibit statistically significantly increased nuclear levels, cellular amounts, or nuclear/cytoplasmic ratio of NF-κB upon LPS stimulation (Figure 4A, B-E). In the absence of LPS, no differences were noted between genotypes in total cellular NF-κB levels or the NF-κB nuclear/cytoplasmic ratio, except for reduced NF-κB nuclear localization in Lmna^-/-^ BMDCs compared to WT BMDCs (Figure 4B-E). In WT-BMDCs with LPS, RelA was predominantly in the nucleus, while in Lmna^-/-^-BMDCs with LPS, RelA was enriched in the cytoplasm, indicating significant differences in nuclear NF-κB levels and nuclear/cytoplasmic ratio (Figure 4A). Additionally, there was a correlation (Pearson two-tailed, P<0.0001) between lamin A/C and NF-κB nuclear levels in both LPS-unmatured and LPS-matured WT cells (Figure 4F). These findings suggest that lamin A/C potentially promotes the nuclear presence of NF-κB in WT-BMDCs, highlighting its role in modulating NF-κB gene-dependent transcription.

**Figure 4.**
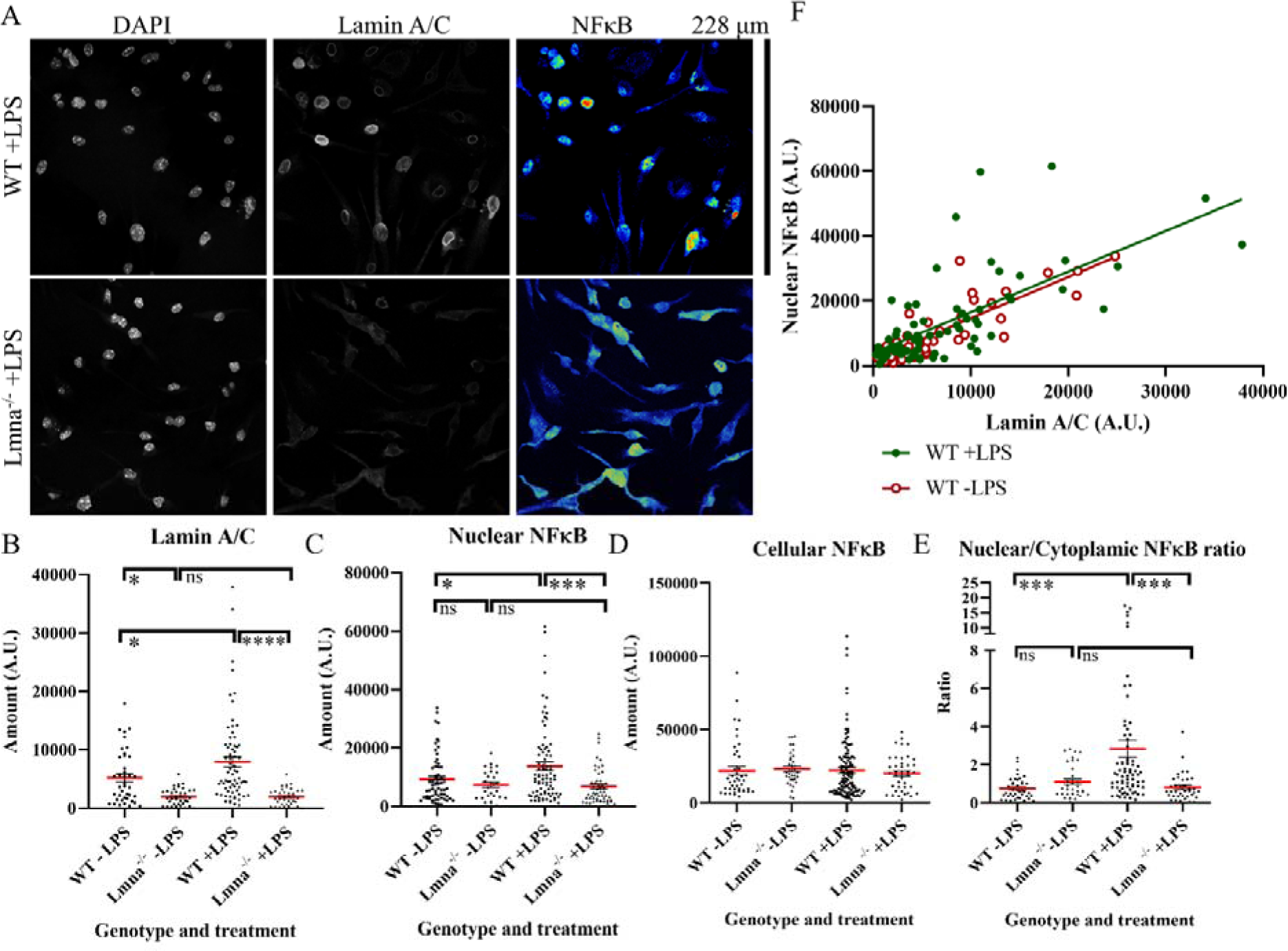
Lamin A/C in GM-CSF-BMDCs favors NF-κB presence inside the nucleus. WT or Lmna^-/-^ GM-CSF-BMDCs, either treated or untreated with LPS for 24 hours, were stained with anti-lamin A/C and anti-NF-κB antibodies along with DAPI. (A) Representative micrographs of LPS-treated WT or Lmna^-/-^ GM-CSF-BMDCs. Quantification of (B) lamin A/C levels, (C) nuclear NF-κB levels, (D) cellular NF-κB levels, and (E) nuclear/cytoplasmic NF-κB ratio. Data are presented as means ± SEM from two independently conducted experiments and were analyzed by one-way ANOVA with Bonferroni’s multiple comparison test (*P < 0.05; ***P < 0.001; ns = not significant; A.U.=Arbitrary units). (F) Correlation between nuclear NF-κB levels and lamin A/C levels in WT GM-CSF-BMDCs, either treated or untreated with LPS. Each symbol in the graphs represents an individual cell (n ≥ 20). The analysis was conducted using linear regression and correlation

### Lamin A/C in BMDCs enhances the expression of NF-**κ**B-related genes

As lamin A/C in BMDCs appeared to promote the presence of NF-κB in the nucleus, we conducted qPCR analysis of genes whose transcription is modulated by this transcription factor. mRNA from WT- and Lmna^-/-^-BMDCs stimulated with LPS for 24 h was extracted, and NF-κB-related gene expression patterns were validated by RT-qPCR (Figure 5). Lmna^-/-^-BMDCs expressed lower levels of proinflammatory molecules such as IL-1α, IL-2, or IFNβ, as well as the anti-inflammatory cytokine IL-10. NF-κB regulates the transcription of genes involved in other functions, including Ccr7 associated with DC migration towards LN, Cxcr2 important for maintaining the fate of normal hematopoietic stem/progenitor cells, Icam1 that encodes an adhesion molecule, and Mip1α, a crucial molecule produced by macrophages and monocytes after bacterial endotoxin stimulation. Expression of these genes was statistically reduced in Lmna^-/-^-BMDCs compared to WT cells.

**Figure 5.**
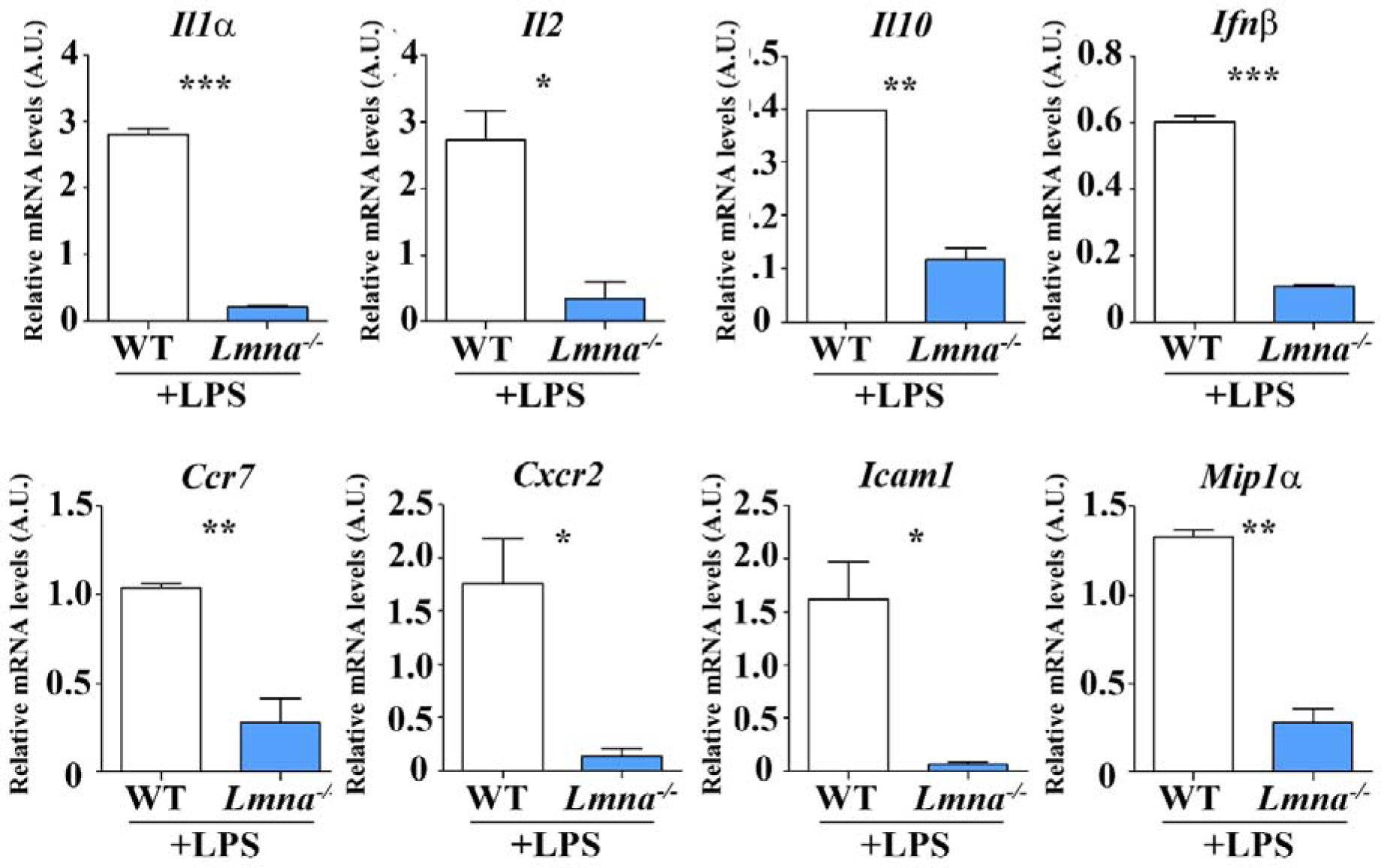
NF-κB-dependent gene expression profiling in WT and Lmna^-/-^-GM-CSF BMDCs stimulated with LPS for 24 Hours. mRNA expression levels of NF-κB-dependent genes in WT- and Lmna^-/-^-BMDCs. The analysis involved RT-qPCR of specified NF-κB-dependent genes in BMDCs stimulated with LPS for 24 hours (n = 3, from 3 independent experiments). The presented data represents means ± SEM and were analyzed by unpaired Student’s t-test. Significance levels are indicated by asterisks (*P < 0.05; **P < 0.01; ***P < 0.001). Housekeeping genes *Ywhaz*, β*-actin*, and *18s* were used for normalization.

These results suggest that lamin A/C may regulate the production of various pro-inflammatory cytokines and other immune molecules, possibly through the modulation of NF-κB localization, among other processes.

### Lamin A/C shapes chromatin accessibility in BMDCs, modulating antiviral response

Lamin A/C is a well-known epigenetic regulator [29,49], yet its influence on transcriptional modulation in DCs during pathogenic infections remains elusive. We aimed to investigate whether alterations in the epigenetic profile of Lmna^-/-^-BMDCs show a distinct innate phenotype in response to LPS stimulation compared to WT-BMDCs. Using ATAC-seq, we examined changes in chromatin accessibility. Isolated LPS-treated WT- and Lmna^-/-^-BMDCs were incubated with transposase 5 (Tn5) to identify regions undergoing active chromatin remodeling. Our analysis identified approximately 21,000 regions with differential accessibility between the conditions (false discovery rate, FDR < 0.05; 2,703 with an abs(log2FC) > 1) (Figure 6A). Functional enrichment analyses of genes associated with differentially accessible sites revealed biological processes compatible with DC and lamin A/C functionalities (Figure 6B, Supplementary Figure 1). Using Reactome pathway analysis via The Database for Annotation, Visualization, and Integrated Discovery (DAVID, NIH), these biological processes corresponded to established lamin A/C functions, such as signal transduction [24], gene expression [50], DNA repair [51], MAPK family signaling cascades, MAP kinase activation [52,53], chromatin-modifying enzymes [28], and chromatin organization [54] (Figure 6B). Additionally, other processes align with crucial functions associated with DCs in viral responses, including antigen processing or presentation, signaling related to TLRs, including TLR4 and 2, and other cytokines (Figure 6B) [55]. Association with immunity were observed using the Gotermmapper tool, as affected genes encompassed categories such as immune system processes, inflammatory responses, and defense response to other organisms (Supplementary Figure 1). Furthermore, several processes are crucial for DC functions in defense responses against other organisms, encompassing signaling, cell differentiation, vesicle-mediated trafficking, cell motility, cell adhesion, membrane organization, autophagy, cell polarity, and lysosome organization (Supplementary Figure 1). Moreover, with this secondary analysis tool, we confirmed that some of these processes directly align with the known functions of lamin A/C [24], including DNA repair, replication, and transcription, signaling, nucleocytoplasmic transport, cell differentiation, cytoskeleton organization, microtubule-based movement, and chromatin organization (Supplementary Figure 1). Hence, these functional enrichment analyses suggest that lamin A/C in DCs, following LPS treatment, potentially primes DCs for a distinct antiviral response. Analyzing these results in depth, we observed that the majority of differential peaks were located in intronic or intergenic regions, spanning 1 to 100 kb from transcription starting sites, indicating their association with regulatory regions. In WT samples, 2840 genes contain peaks that were significantly higher, with a P-value less than 0.05 compared to the other genotype while Lmna^-/-^ samples had 44 genes with higher peaks (Figure 6A). Among these, 2152 (75%) were in promoters, 197 (6.9%) in CpG Islands, and 177 (6.2%) in 5’ UTR regions in the WT, whereas 3 (7.0%), 3 (6.8%), and 2 (4.5%), respectively, were observed in these locations in the Lmna^-/-^ samples. This data suggests reduced chromatin accessibility in Lmna^-/-^-BMDCs compared to WT-BMDCs in regions crucial for transcriptional regulation.

**Figure 6.**
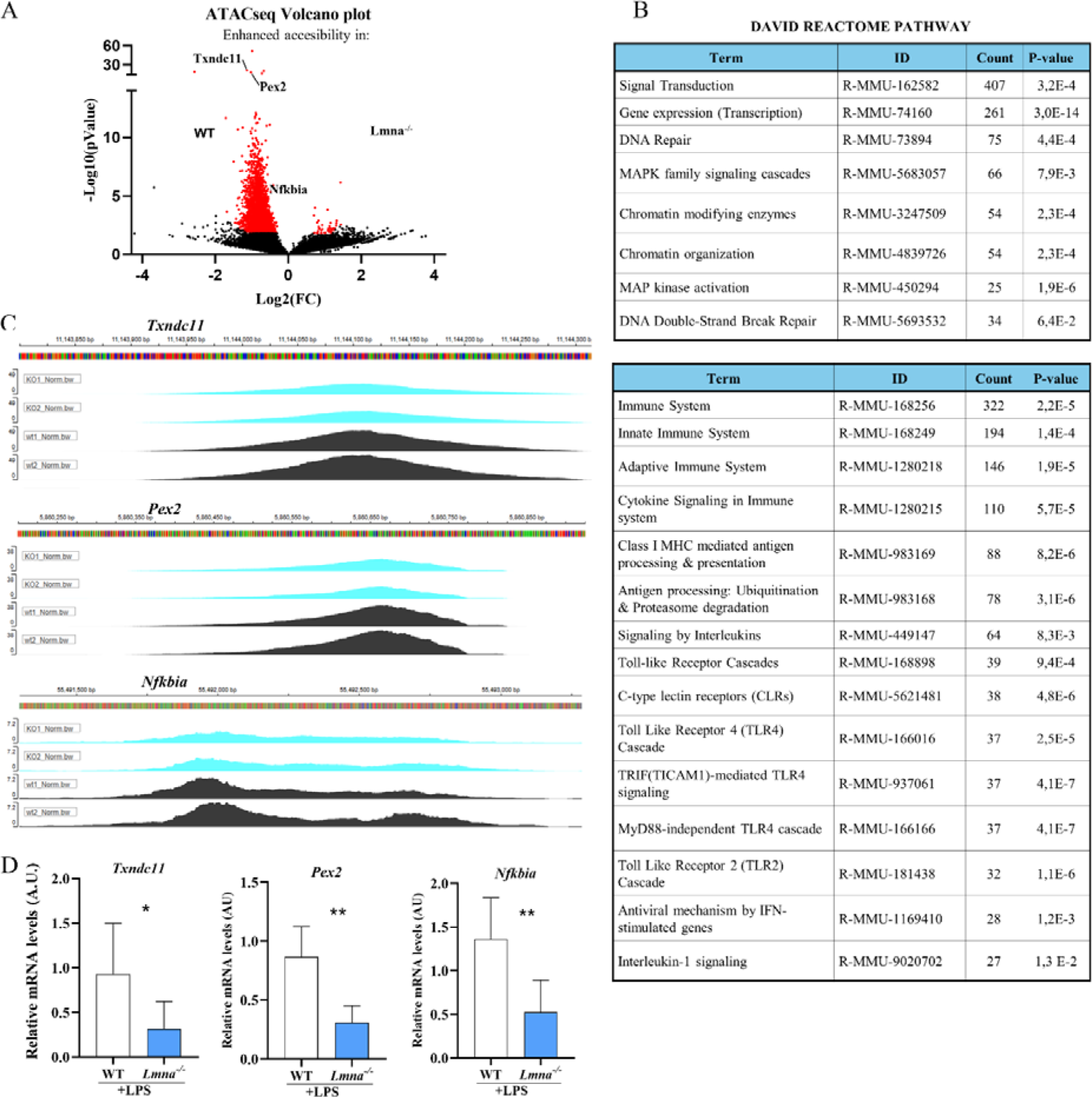
Differential epigenomic signature in WT and Lmna^-/-^ GM-CSF BMDCs upon LPS stimulation. (A) Visualization of the results from the differential accessibility analysis using ATAC-seq data. Volcano plot representing differential accessibility analysis results for ATAC-seq data. Differentially accessible sites [with FDR<0.05 and abs(log2FC)>1] are symbolized as red dots. (B) Enriched biological processes (GO) terms detected with DAVID Reactome pathway were associated with differentially accessible site proximal genes. Only the most enriched terms related to immune system, DCs, or lamin A/C functions are presented, sorted by the number of genes detected or P-value. (C) Peak representation of DNA accessibility in WT GM-CSF BMDCs (dark) compared to Lmna^-/-^ BMDCs (soft) within regions proximal to the locus of the indicated genes. The height of the peak reflects the counts per million from the ATAC-seq data (n = 2). (D) mRNA expression levels measured by RT-qPCR analysis for the indicated genes in WT and Lmna^-/-^ BMDCs stimulated with LPS for 24 hours. The data, obtained from three independent experiments with three samples each, were analyzed using unpaired Student’s t-test. Housekeeping gene *18s* was used for normalization. Data are presented as means ± SEM. (*P < 0.05; **P < 0.01; ***P < 0.001; ns, not significant).

To validate the functionality of the accessibility of some of the regulatory regions of these genes, we confirmed that genes with higher chromatin accessibility in WT-BMDCs, such as Tnxdc11, Pex2 and Nfkbia (Figure 6A,C), were up-regulated at the mRNA expression level (Figure 6D). These differences in DNA accessibility between WT- and Lmna^-/-^-BMDCs may indicate a lamin A/C-dependent epigenetic ability to modulate DC functions in the immune response against viruses.

## Discussion

This study demonstrates that downregulation of lamin A/C in LysM-expressing myeloid cells suppresses their ability to generate effector CD8 CTL and Th1 responses *in vivo*, as well as a Th1 response in naïve T cells stimulated *in vitro* with BMDCs. This effect is mediated by changes in immune synapse formation, nuclear presence of NF-κB, and the epigenetic signature, ultimately leading to the downregulation of proinflammatory gene expression.

The clearance of VACV in peripheral infection involves both CD8 CTL and Th1 cells through various mechanisms [56]. Approximately 7 days after VACV infection [57], there is an expansion of VACV-specific CD8 T cells peaking 10 days post-infection [33]. Additionally, the primary VACV-specific CD4 T cell response peaks at around 14 days post-inoculation, characterized by highly activated CD4 T cells producing IFNγ [58]. These IFNγ-producing CD4 T cells play a role not only as cytotoxic cells [59,60] but also as helpers for CD8 CTLs, supporting the maintenance of the primary CD8 CTL response in an MHCII-dependent manner [33].

In our study, there is a temporal correlation between the reduced VACV eradication observed at 14 days in Lmna^-/-^ mice, reduced Th1 cell production at both 6 and 14 days, and decreased CD8 CTL production only at 14 days in lamin A/C-deficient mice upon s.s. These findings emphasize the importance of lamin A/C in the myeloid compartment for generating a Th1 and CD8 CTL response against the virus. They suggest that the increased viral accumulation in Lmna^-/-^ mice at 14 days may result from reduced viral clearance due to the diminished CD8 CTL and Th1 response. It is not yet clear whether this reduced CD8 CTL response at 14 days is a direct result of the interaction with lamin A/C-deficient APCs or an indirect effect of the reduced Th1 response observed at 6 days, which may itself be a consequence of the absence of lamin A/C in APCs. Despite our results, the significance of CD4 T cells in the primary CD8 T cell response to VACV infection remains somewhat controversial [61], with some studies showing dependence [62–64], while others do not [22,65,66]. This variability in results may be linked to the route of infection and viral dose [33,34]. Our results confirm that the route of VACV administration affects the primary CD4 and CD8 T cell responses. Since the absence of lamin A/C in APCs leads to differences in the CD8 CTL and Th1 responses depending on the administration route.

We were interested in establishing the cellular and molecular mechanisms that could be regulated by lamin A/C and thus modulate the functions of DCs in the antiviral response. Our results showed that lamin A/C absence in BMDCs stimulated with VACV lysate significantly reduced their ability to promote activation, proliferation, and differentiation into a Th1 phenotype. VACV can be recognized by TLR2[37,38] and TLR4[39,40] and indeed, the absence of lamin A/C in BMDCs matured with a TLR2 ligand inhibits their ability to promote activation and differentiation towards a Th1 phenotype, particularly at later times, without affecting early activation levels or T cell proliferation under the conditions studied. Additionally, after maturation of BMDCs with LPS, BMDCs have a reduced capacity to promote both early and late activation, as well as proliferation and differentiation into Th1 cells in co-cultures where cells lacking lamin A/C participate. This reduced effect observed after maturation of BMDCs with a TLR2 ligand, compared to the effects observed with VACV lysate or LPS, may be due to the greater stimulation of different receptors in the case of VACV and to a greater importance of lamin A/C in the pathways stimulated by LPS in the latter case. This led us to continue our research using LPS-mediated maturation of the TLR4 signaling pathway in the following experiments.

Upon recognition of LPS by TLR4, DCs undergo a complex developmental program known as maturation, resulting in significant modifications in both structure and function [67]. This maturation enhances DC migration and upregulates the expression of costimulatory molecules, thereby increasing the capacity of DCs to present exogenous antigens [67].

*In vivo*, once matured and after incorporation, processing, and presentation on the membrane in the MHCII context of VACV antigens, DCs migrate to a lymph node where they will specifically stimulate T cells, including CD4 T cells, through the formation of immune synapses [67]. The observed reduced capacity to generate Th1 cells could be related to a decreased formation of immune synapses by Lmna^-/-^ DCs [26]. Our videomicroscopy experiments confirmed this possibility but without changes in MHCII or costimulatory molecules.

TLR4 signaling promotes NF-κB-related genes expression [45]. NF-κB plays a crucial role in regulating DC development, survival, and cytokine production [46]. Previous studies have indicated a role for lamin A/C in enhancing the nuclear translocation of RelA/p65 and subsequently increasing NF-κB activity, leading to elevated inflammatory gene expression in macrophages [48]. We observed that upon LPS addition, in WT-BMDCs, RelA was predominantly in the nucleus, while in Lmna^-/-^-BMDCs, RelA was enriched in the cytoplasm. These findings suggest that the absence of lamin A/C ameliorates the nuclear presence of NF-κB in the nucleus, potentially reducing NF-κB gene-dependent transcription.

The investigation into how lamin A/C triggers NF-κB translocation remains unresolved. Some researchers propose that lamin A/C activates IKKβ, leading to the phosphorylation and subsequent reduction of IκBα protein levels, thereby facilitating NF-κB nuclear translocation and transcriptional activation [48]. Another hypothesis suggests that the accumulation of prelamin A at the nuclear lamina activates the NF-κB pathway via NEMO-dependent signaling or inflammasome formation [68,69]. Additionally, lamin A/C may modulate actin dynamics, influencing the nuclear translocation and downstream signaling of transcription factors like megakaryoblastic leukemia 1 (MLK1) [70]. Conversely, downregulation of lamin A/C could potentially diminish Rel A retention in the nucleus, resulting in NF-κB hypoactivation and the attenuation of inflammatory cascades. Another potential mechanism involves the regulation of nuclear NF-κB transcriptional activity through chromatin structure reorganization. Recent studies have illustrated changes in 3D genome organization during immune activation, wherein immune gene loci are released from the nuclear periphery to facilitate proper immune response activation [71]. Reduced lamin A/C levels may alter 3D chromatin organization or the interaction of the genome with the nuclear periphery, leading to dysregulated expression of inflammatory genes. To further explore this possibility, we investigated whether the absence of lamin A/C in DCs could modify the epigenetic profile of these cells. Lamin A/C is a well-known epigenetic regulator [49], yet its impact on transcription modulation in DCs during pathogen infections remains unclear. We aimed to investigate whether alterations in the transcriptomic profile of Lmna^-/-^-BMDCs indicate a distinct innate phenotype in response to LPS stimulation compared to LPS-treated WT BMDCs. Using ATAC-seq, we assessed changes in chromatin accessibility. Functional enrichment analyses of genes associated with differentially accessible sites align directly with the known functions of lamin A/C in other contexts [24], and suggest the importance of lamin A/C on these critical DC functions in the context of viral infection. Also, these findings expand our understanding of the role of lamin A/C in these functions within the context of immune system cells. This functional enrichment analysis suggests that lamin A/C in DCs, following LPS treatment, primes DCs for an enhanced antiviral response. Confirming this, the affected genes are annotated in general categories such as immune system processes, inflammatory responses, and defense responses to other organisms.

This ATAC-seq results also showed that the majority of differential peaks were situated in regulatory regions. In WT samples, a higher number of peaks were significantly higher, with a P-value smaller than 0.05 compared to Lmna^-/-^ samples. This suggests reduced chromatin accessibility in Lmna^-/-^-BMDCs compared to WT BMDCs in regions crucial for transcriptional regulation. This is in accordance with the increased general capacity of activation of T cells as observed in previous figures.

In summary, these results underscore the critical role of lamin A/C in APCs, particularly moDCs and BMDCs, during the immune response against viral infections. They emphasize the importance of lamin A/C in optimal T cell activation, differentiation, and antiviral immune responses, providing insights into potential therapeutic strategies targeting APCs in infectious diseases.

## Methods

### Mice

C57BL/6-Tg (TcraTcrb)425Cbn/J mice (CD45.1 OT-II mice) expressing a TCR specific for the OVA peptide (amino acid residues 323 to 339) in the context of I-Ab were obtained from the Jackson Laboratory (stock number 004194). The LysM-cre Lmna^flox/flox^ mouse (referred as LysM-Lmna^-/-^) was generated by crossing C57BL/6 LysM-cre^+/-^ mice [72] with C57BL/6 Lmna^flox/flox^ mice [73]. C57BL/6 Lmna^flox/flox^ littermates were used as WT control mice. C57BL/6-CD45.2 and C57BL/6-CD45.1 WT mice were used as recipients for adoptive transfer. Lmna^flox/flox^ mice were generously provided by Y. Zheng [73]. All mice were bred at the CNIC under specific pathogen-free conditions. Experiments were performed using mice that were matched for both sex and age (6-12 weeks). The mice were maintained on a standard 12-hour light/dark cycle (with the light period from 7 a.m. to 7 p.m.), and they had access to food and water ad libitum. The Animal Care and Ethics Committee of the CNIC, UAM, and the regional authorities approved all experimental procedures.

### Antibodi22es and Reagents

Antibodies against: CD4 (Clone RM4-5, GK1.5) -v450, -APC and PE; CD25 (PC61.5) -APC; CD69 (H1.2F3)-FITC; CD28 (37.51); CD3 (145-2C11); CD45.1 (A20)-Pe-Cy7, -v450, -FITC; CD45.2 (104) -v450, -redFluor710; CD11b (M1/70)-v450; CD86 GL-1 PE; IFNγ XMG1.2 FITC, -APC; Foxp3 (3G3)-FITC; MHC-II (I-A/I-E, M5/115.15.2)-APC; GR1 (RB6-8C5)-Biotin; CD80 (16-10A1)-Biotin; B220 (RA3-6B2)-Biotin; F4/80 (BM8.1)-Biotin; CD11b (M1/70)-Biotin; CD19 (1D3)-Biotin; CD25 (PC61.5)-Biotin from Tonbo. CD11c (N418)-FITC from BioLegend. Lamin A/C (4C11)-Alexa488; NF-κB (D14E12-XP) from Cell Signaling. IgM (R6-60.2)-Biotin; MHC-II (2G9)-Biotin; CD49b (DX5)-Biotin; Streptavidin-PE from BD Bioscience. Lipopolysaccharide (LPS) (from Sigma-Aldrich, Catalog number L2630, final concentration 20 ng/ml); Palmitoyl-3-CysSerLys-4 (Pam3CSK4) (InvivoGen, tlrl-pms, 100 ng/ml); 7-amino-4 chloromethylcoumarin (CMAC) CellTrakerTM (Invitrogen, C2110, 0.1 µM); Vybrant CFDA SE Cell Tracer Kit-Carboxyfluorescein diacetate succinimidyl ester (CFSE) (Invitrogen, V12883, 5 µM); CellTrace Violet (CTV) (Invitrogen, C34557 5 µM); Cell Stimulation Cocktail (500X) (Tonbo, TNB-4975-UL100, 1X); 4’,6-Diamidino-2-Phenylindole, Dihydrochloride (DAPI) (Invitrogen, D1306, 1:10000); Vybrant DiL Cell-Labeling Solution (DiL tracer) (Invitrogen, V22885, 1:1000); Mouse GM-CSF (PeproTech, 315-02, 20 ng/ml); OVA-OT-IIp(OVA323-339, ISQAVHAAHAEINEAGR) (GenScript, 10 μg/ml); Fluorescence Mounting Medium (Dako Omnis) with 500 µg/l DAPI (Agilent, GM30411-2, 30 µl/sample)

### T cell isolation

Spleen and LN cell suspensions from CD45.1 OT-II mice were obtained by homogenization through a 70 μm cell strainer (Falcon). In spleen suspensions, red blood cells (RBC) were lysed using eBioscience Red Blood Cell Lysis Buffer (Invitrogen) for 5 min at RT. Naïve CD4 T cells were isolated using the EasySep Kit or negative selection with labeled antibodies (anti GR-1, B220, F4/80, CD8α, CD11b, CD19, CD25, IgM, MHC-II and CD49b) and streptavidin-bound magnetic microbeads (Miltenyi Biotec). Isolated CD4 T cells were activated with anti-CD3 and anti-CD28 antibodies if needed.

### Generation of Bone Marrow-Derived DCs

Bone marrow-derived DCs (BMDCs) were generated as described by [74]. Briefly, femur and tibiae were extracted from 6-12 weeks old WT and LysM-Lmna^-/-^ mice. After removing surrounding muscle tissue, both bone ends were cut, and marrow was flushed with PBS using a 25G syringe. RBCs were lysed with eBioscience Red Blood Cell Lysis Buffer. The bone marrow cell suspension was cultured on non-treated 150-mm Petri dishes at a concentration of 0.5×10^6^ cells/ml in complete RPMI-1640 with 20 ng/ml rGM-CSF (PeproTech). On day 3, attached cells were removed, and the remaining cells were cultured until days 7-9. Maturation was induced based on the experiment, either with an overnight incubation with TLR4 activator Lipopolysaccharide (LPS), 4-hour treatment with TLR2 activator Pam3CsK4, or with irradiated VACV extract (stock 10^7^ p.f.u, use 1:1000) at 37 °C. On days 7-9, BMDCs were collected and used for subsequent experiments.

### BMDCs and CD4 T cells *in vitro* co-cultures

Naïve CD4 T cells were isolated from CD45.1 OT-II mice, and GM-CSF-BMDCs were matured with LPS (overnight), Pam3CsK4 (4 hours), or irradiated VACV extract (4 hours), followed by pulsing with 10 µg/ml OVA323–339 peptide (OVA-OT-IIp) for at least 30 minutes at 37 °C. Simultaneously, naïve CD4 T cells were labeled with CellTrace Violet following the manufacturer’s protocol (Life Technologies). Co-cultures of BMDCs and CD4 T cells were conducted with a 1:2 ratio, respectively, in 96-well U-bottom plates. CTV-labeled CD4 OT-II T cells cultured alone served as a negative control, and CD4 OT-II T cells stimulated with plate-bound anti-CD3 (5 μg/ml; Tonbo) and soluble anti-CD28 (1 μg/ml; Tonbo) antibodies served as positive controls. Activation, proliferation, and differentiation of naïve CD4 OT-II T cells in response to OVA323–339 peptide-primed BMDCs subsets were assessed by flow cytometry, monitoring specific markers (CD25 and CD69 for activation, IFNγ for Th1 differentiation, and IFNγ-producing CD8 T cells) and the reduction in CTV intensity at 24 hours, 48 hours, and 6 days.

### BMDCs and CD4 T cells *in vivo* co-cultures

For adoptive transfer experiments, CD45.1 OT-II recipient mice received a subcutaneous injection (s.c.) in footpad with 2×10^6^ WT- or Lmna^-/-^-BMDCs matured with LPS and pulsed with OVA323–339 cognate OT-IIp. Six days after BMDC injection, draining popliteal lymph nodes (pLN) were extracted, and Th1 differentiation was assessed by flow cytometry.

### Flow Cytometry

Single-cell suspensions were transferred to a 96-V-well plate and pre-incubated with a purified anti-CD16/32 antibody (FcBlock, Tonbo) for 10 min on ice before immunostaining. Cells were then stained with the appropriate surface marker antibodies, on ice for at least 30 min according to the manufacturer’s instructions. Finally, a viability marker was added to the labeled cell suspension. To assess IFNγ production intracellularly, CD4 T cells were stimulated with Cell Stimulation Cocktail (500X) (Tonbo) for 4 h at 37 °C to allow intracellular cytokine accumulation. After surface staining, samples were fixed and permeabilized with a Cytofix/Cytoperm kit (BD Biosciences) for 20 min at 4 LC. The solution was washed out with Perm/Wash™ Buffer (BD Biosciences), and then, cells were intracellularly stained with anti–IFNγ for 1 h at RT. For Foxp3 staining, the mouse Foxp3 Buffer set (BD Pharmigen) was used following the manufacturer’s protocol. For lamin A/C intranuclear staining, cells were fixed with 4 % paraformaldehyde (PFA) and permeabilized with 0.5 % Triton X-100/PBS solution for 5 min at RT. Anti-mouse lamin A/C antibody was then incubated for 1 h at RT. Data were acquired on LSRFortessa or FACSCanto flow cytometers (BD Biosciences) and analyzed with FlowJo V10.4.2 software (Treestar Inc.).

### Time-Lapse Microscopy

For analysis of T cell-BMDC cognate interactions, WT or Lmna^-/-^-BMDCs matured with LPS for 24 h and pulsed with OVA323–339 cognate OVA-OT-IIp were mixed with CD4 OT-II T cells (1:1). To observe live T cell-BMDC interactions, BMDCs labeled with DiL tracer (Vybrant™ DiI Cell-Labeling Solution, Invitrogen) were suspended in Hanks balanced salt solution (HBSS) (Lonza) with 2% FBS and 20 mM HEPES (Hyclone) and plated onto poly-L-Lysine (PLL; 20 µg/ml, Sigma-Aldrich)-coated 35-mm-diameter culture dishes (MatTek) for at least 15 min at 37 °C. After BMDCs adhered to the PLL coating, CD4 T cells previously labeled with CMAC cell tracer (7-amino-4-chloromethylcoumarin, CellTracker Blue CMAC Dye, Invitrogen) were added to the microscope (Nikon ECLIPSE Ti-TimeLapse microscopy), covered by an acrylic box for acquisition at 37 °C and 5% CO2. T cell-BMDC interactions were recorded for 150 min.

### Fluorescence Confocal Microscopy

For immunostaining, WT- or Lmna^-/-^-BMDCs were treated or not with LPS for 24 h and then plated onto PLL-coated coverslips (20 μg/ml) for 30 min at 37 °C. Subsequently, cells were fixed in 4% PFA, permeabilized in 0.5% Triton X-100/PBS solution for 5 min, and stained with appropriate antibodies: anti-NF-κB p65 and Alexa488-conjugated anti-mouse lamin A/C antibodies (both from Cell Signaling) and DAPI (Invitrogen) for 1 h at RT. Finally, samples were mounted in Prolong antifade medium (Invitrogen). Confocal images were acquired using a Leica gated STED-3X-WLL SP8 microscope under the same conditions, including identical exposure times and light intensity. Image analysis was performed using ImageJ Fiji software and Imaris software (Bitplane).

### Reverse transcription-quantitative PCR

Total RNA from cultured WT or Lmna*^-/-^*-BMDCs (treated with LPS for 24 h) was isolated with TRI Reagent solution (Invitrogen) and isopropanol precipitation. RNA purity and concentration were assessed from the ratio of absorbance at 260 and 280 nm in a Nanodrop-1000 Spectrophotometer (Thermo Scientific). Total RNA (0.5 to 2 μg) was reverse transcribed to complementary DNA (cDNA) with the High-Capacity cDNA Reverse Transcription Kit with RNase Inhibitor (Applied Biosystems). Quantitative PCR was performed using the PCR Power SYBR Green PCR Master Mix (Applied Biosystems) in a B7900-FAST-384 Sequence Detection System (Applied Biosystems), with technical triplicates. Expression levels of target genes were normalized to at least 2 housekeeping genes: β*-actin*, *Ywhaz* (Tyrosine 3-Monooxygenase/Tryptophan 5-Monooxygenase Activation Protein Zeta), or *18s.* Gene-specific primers used, and their sequences are listed in Table 1.

**Table 1.**
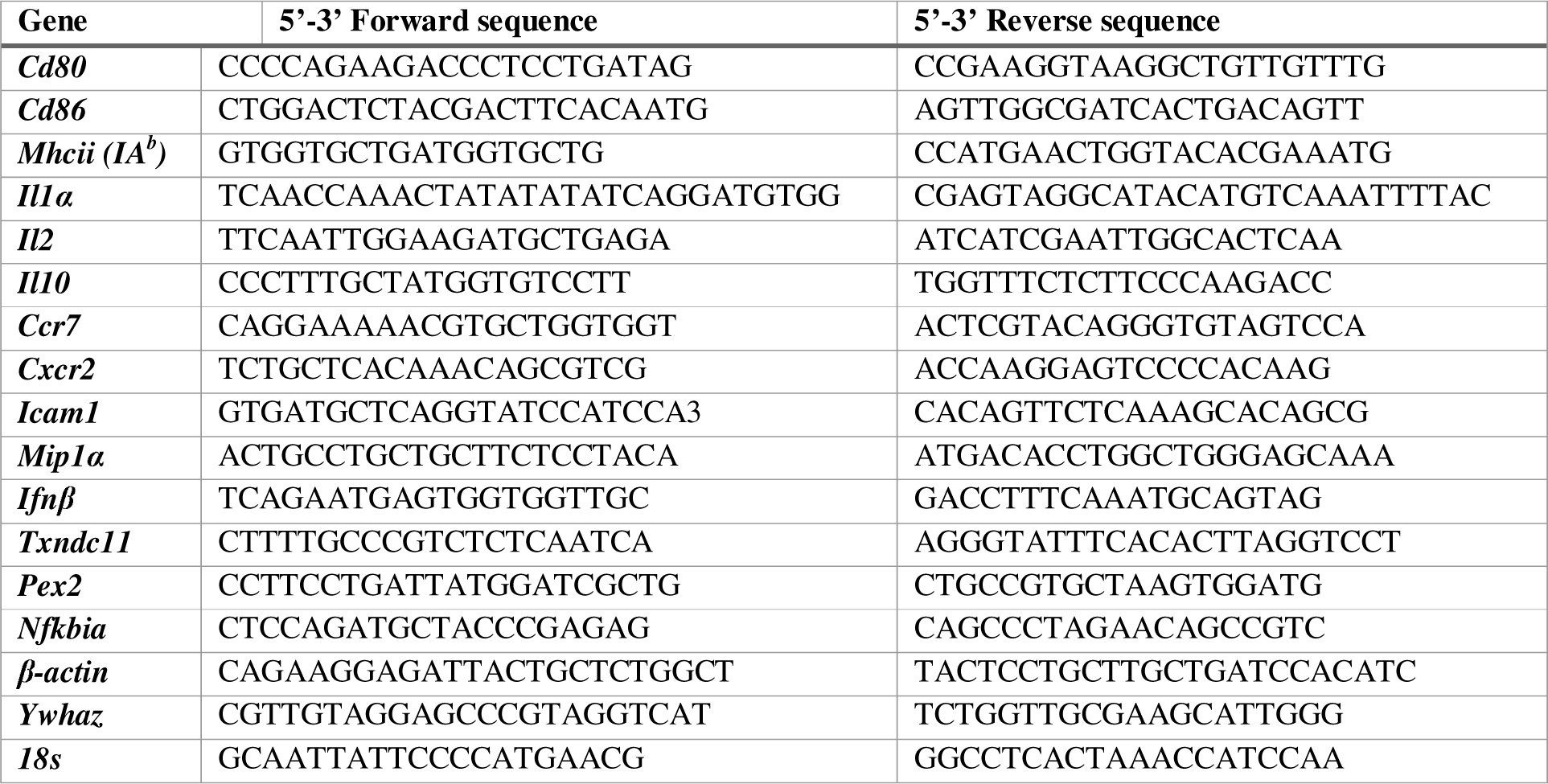
List of the primers used for RT-qPCR.

### *Vaccinia virus* (VACV) *in vivo* infection and viral titration

The VACV strain Western Reserve (WR, ATCC number VR-1354) was kindly provided by Jonathan W. Yewdell and Jack R. Bennink (NIH, Bethesda, Maryland, USA). VACV was propagated in African Green Monkey Kidney Fibroblast Cells (CV-1) and used as clarified sonicated cell extracts. WT or OVA-expressing VACV were administered by intraperitoneal (i.p.) injection, subcutaneous (s.c.) inoculation in the footpad [75] or by skin scarification (s.s.) in the tail of WT and LysM-Lmna^-/-^ mice. 1×105 or 1×106 plaque-forming units (p.f.u.) were used depending on the route of administration. For this last procedure, animals were anesthetized with inhalatory isoflurane, and then, thirty scarifications were made with a 25 G syringe in 1 cm along the base of the tail avoiding bleeding, as previously described [76]. After that, 1 ml of PBS containing 1×106 p.f.u. of VACV-WR was added to the area and allowed to air dry. Only PBS was added to the mock-infected animals. For viral quantification, mice were euthanized, and the tail was aseptically removed, weighed, frozen, and stored at -80 °C until usage. The samples were homogenized with a Tissue Ruptor (QIAGEN, USA) in 0.5 ml DMEM containing 50 U/ml penicillin and 100 μg/ml streptomycin. The homogenates were sonicated for 3 min at 40 % amplitude, freeze-thawed twice (-80 °C/ 37 °C), sonicated again under the same conditions, and then serially diluted in DMEM without FBS. To quantify p.f.u., dilutions were added to monolayers of CV-1 seeded on 24-well plates and incubated at 37 °C and 5 % CO2 for 1 h. Hereafter, 0.5 ml of DMEM containing 0.5 % FBS was added to each well. After 24 h, cells were stained for 5 min with crystal violet solution (0.5 % crystal violet, 10 % ethanol, and 1 % PFA) and washed again. Viral plaques were counted, and data was presented as p.f.u./g [56].

### ATAC-seq

ATAC-seq was performed as described in [77]. A number of 5×10^4^ CD11c^+^ BMDCs were sorted and lysed [10 mM tris-HCl (pH 7.4), 10 mM MgCl2, and 0.1% IGEPAL CA-630] for 15 min at 4°C while centrifuging at 500g. The supernatant was discarded, and nuclei were resuspended in transposase reaction buffer with Tn5 and incubated for 1 hour at 37°C and 7 min at 4°C. Later, 0.1% SDS was added to stop the reaction (5 min, room temperature), and DNA was isolated with Agencourt AMPure XP [Beckman Coulter; solid phase reversible immobilization (SPRI) beads, 2×] and measured with Qubit Kit. For DNA library generation, qPCR with i5 and i7 primers (barcoding) was performed (98°C 2′, 98°C 20L, 63°C 30L, 72°C 1′, and 4°C ∞; 9 cycles). A size cut-off was conducted (0.65× SPRI beads), and DNA was isolated with 1.8× SPRI beads. Last, another qPCR was performed with i5 and i7 primers with 6 to 7 cycles for 0.9 ng/μl or with 5 cycles for more than 0.9 ng/μl (98°C 2′, 98°C 20L, 63°C 30L, 72°C 1′, and 4°C ∞), and amplified DNA was isolated with SPRI beads (2×), and its concentration was measured by Qubit. As control, we used qPCR for *Txndc11*, *Pex2*, *Nfkbia* promoter. An Agilent Pico 6000 bioanalyzer was used to assess peak sizes and percentages. Libraries were sequenced on a HiSeq 2500 (Illumina) to generate 2 × 100–base pair (bp) reads and processed with RTA v1.18.66.3. FastQ files for each sample were obtained using bcl2fastq v2.20.0.422 software (Illumina). Sequencing reads were trimmed off Illumina adapters and transposase sequence using cutadapt 1.16. Then, they were aligned to the mouse reference genome (GRCh38 v91) using bowtie 1.2.2 with a limit of 2000-bp paired distance. The aligned bam files were converted to bw for visualization purposes using bamCoverage from deepTools.

### Bioinformatics analysis

Sequencing reads were preprocessed with a pipeline that used FastQC to assess read quality and Cutadapt to trim sequencing reads, eliminating Illumina adaptor remains, and to discard reads that were shorter than 30 bp. In the case of ATAC-seq, Nextera transposase adapter contaminations were also removed with Cutadapt. Preprocessed ATAC-seq reads were mapped against reference genome mm9 (same assembly than NCBIM37) with a pipeline that used the Burrows-Wheeler Aligner-Maximal Exact Match (BWAMEM) algorithm as aligner, Piccard to mark duplicate alignments, and samtools to eliminate duplicates, chimerism, and sub-optimally multimapped alignments. Only properly paired and mapped reads were kept. Alignments against the mitochondrial genome or chromosomes X and Y were also removed. The final number of read pairs was 10 to 20×10^6^ for any of the samples. Transcription start site (TSS) enrichment values, calculated with HOMER’s annotate Peaks function, had values between 4 and 11 for any of the samples. Once filtered alignments had been obtained, peaks (accessible DNA regions) were called with MACS2, using parameters “--nomodel --shift -100 --extsize 200”, and “-q 0.05” as the FDR cut-off. Around 100,000 peaks were detected for any of the samples. Next, filtered alignments and peaks, in bam and bed formats, respectively, were processed with the R package DiffBind to define a consensus set of 98,731 peaks, to calculate and normalize their coverage in various samples, and to identify differentially accessible regions, using edgeR as analysis method. The fraction of reads in peaks (FRiP) score, as calculated by DiffBind, was around 0.5 for any of the samples. Functional enrichment analyses for the set of differentially accessible regions were performed with the genomic regions enrichment of annotations tools GOTermMapper [78,79] and DAVID [80,81] following developer instructions. Other data manipulations and graphical representations (heatmaps, bars, and scatterplots) were produced with R or Graphpad.

### Statistical analysis

Statistical analyses were performed with GraphPad Prism 8 software. Normality of the data was tested using the Kolmogorov-Smirnov test. The parameters followed a normal distribution, and, therefore they were represented as the distribution of each sample, mean ± SEM (Standard Error of the Means). Parametric tests were used for statistical analysis. Unless otherwise stated, statistical significance was calculated by Unpaired Student’s t-test. When specified, one-way ANOVA or two-way ANOVA were used, with Bonferroni’s post-hoc multiple comparison test applied as appropriate. The significance of differences was calculated as follows: *P < 0.05, **P < 0.01, and ***P < 0.001. Not significant was depicted as n.s.

## Acknowledgments

We thank Hector Sanchez Martinez and Tamara del Castillo for his technical assistance. This work was supported by grants from Ministerio de Ciencia, Innovación y Universidades (MCNU from Spain (grant number RTI2018-097504-B-I00; PID2021-125415OB-I00), Ministerio de Economía y Competitividad (grant number PID-2020-120412RB-I00) and La Caixa Health Research Grant (grant number LCF/PR/HR23/52430018). Instituto de Salud Carlos III (ISCIII) (grant number PI20/00306) with co-funding from the European Regional Development Fund (ERDF) “A way to build Europe”. The CNIC is supported by the ISCIII, the Ministerio de Ciencia, Innovación y Universidades (MCNU), and the Pro CNIC Foundation. B.H.-F. and R.G.-B. by the UAM and the MCNU FPU program (FPU18/00895, FPU19/01774); A.S. by Universidad Francisco de Vitoria; and H.S-M. by the Comunidad de Madrid YEI program (PEJ-2020-TL/BMD-17604).

## Author Contributions

B.H.F; R.G.B; M.O.Z: Data curation, Formal Analysis, Investigation, Methodology. Software, Validation, Visualization. S.I; F.S.M: Conceptualization, Funding acquisition, Investigation, Project administration, Writing-review & editing. A.Q; A.D: Funding acquisition, Investigation, Data curation, Formal Analysis, Writing-review & editing. E.V: Data curation, Formal Analysis. A.S: Conceptualization, Formal Analysis, Investigation, Methodology, Writing-original draft, Writing-review & editing. V.Z: Data curation, Formal Analysis, Investigation, Methodology. S.M.A: Conceptualization, Investigation, Funding acquisition, Investigation, Project administration, Resources, Supervision, Validation, Visualization, Writing-review & editing. J.M.G.G. Conceptualization, Data curation, Formal Analysis, Investigation, Funding acquisition, Investigation, Project administration, Resources, Software, Supervision, Validation, Visualization, Writing-original draft, Writing-review & editing.

---

**Supplementary Figure 1.**
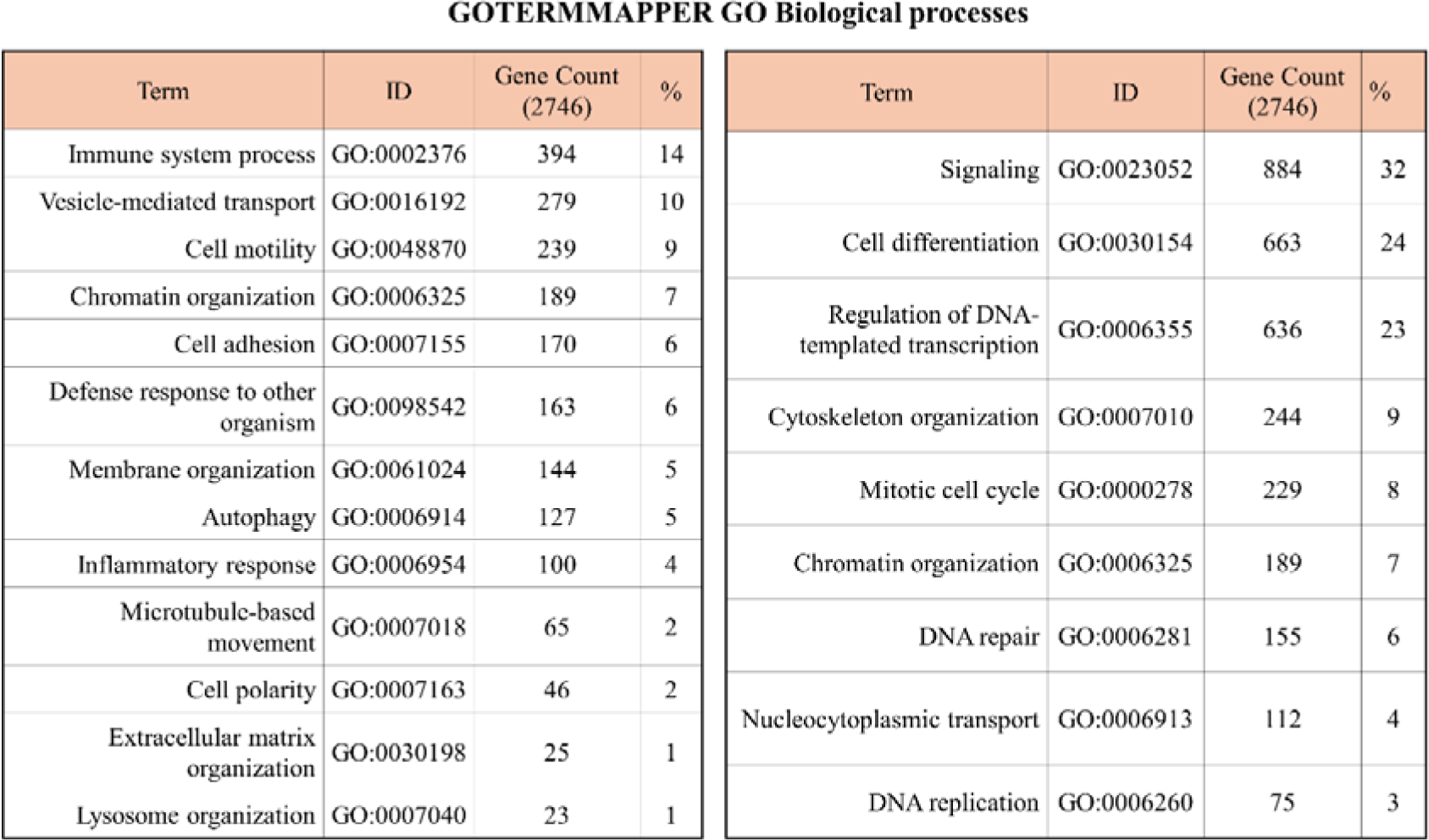
Differential biological processes (GO) terms in WT and Lmna-/- GM-CSF BMDCs upon LPS stimulation. Enriched biological processes GO terms, detected with GOTERMMAPPER were associated with differentially accessible site proximal genes. Only the most enriched terms related to immune system, DCs, or lamin A/C functions are presented, sorted by the number of genes detected or P-value.

